# E-bike riding: A metabolic evaluation in the context of exercise intensity domains

**DOI:** 10.1101/2024.11.21.624713

**Authors:** Alberto Bonardi, Francesco Negro, Danilo Iannetta

## Abstract

**Introduction:** E-bikes are being promoted as a mode of transportation that can aid with meeting current physical activity guidelines. This study evaluated exertional intensity associated with E-Biking within the exercise intensity domain framework. We hypothesized that exertional intensity of E-bikes is insufficient to evoke a metabolic demand associated with the minimum intensity needed to improve cardiorespiratory fitness.

**Methods:** Forty-four participants (22 females) of varying activity levels completed two experimental sessions. The first session involved a lab-based ramp-incremental exercise test to determine V̇O_2max_, gas exchange threshold (GET), and respiratory compensation point (RCP). The second session involved the completion of two outdoor rides on the same bike equipped with an electrical motor E-bike. The first ride was completed without electrical assistance, while the second was. The power output (PO) and speed were set at an intensity corresponding to ∼10% above GET.

**Results:** As expected, using E-bike assistance resulted in a lower metabolic demand, falling well below GET. While the absolute power output (PO) was different between sexes, relative heart rate (HR) and relative PO were similar between the rides without and with assistance. This suggests that when riding an E-bike, the internal load is similar between sexes.

**Conclusion:** Despite E-bikes facilitate a more active lifestyle and help to reduce the emission of pollutants, when interpreted within the context of the exercise intensity domain schema, their associated exertional intensity is likely insufficient to confidently elicit cardiorespiratory benefits.

## Introduction

Sedentarism and physical inactivity remain a public health concern, worsening quality of life and contributing to a substantial proportion of all-cause mortality globally (Park et al., 2020; Patterson et al., 2018). Currently, one in four adults and three in four adolescents do not meet the physical activity guidelines of at least 150 minutes of moderate- (3-6 METs) or 75 minutes of vigorous-intensity (6-9 METs) physical activity per week that is recommended to maintain health (Bull et al., 2020).

In recent years, the use of electric bikes (E-bikes) has spread considerably, being promoted as a mode of transportation to not only reduce traffic congestion and pollution (Bourne et al., 2020; Johansson et al., 2017; Li et al., 2023; Sundfør et al., 2020) but also to help individuals of all ages to adopt a more active lifestyle (Bourne et al., 2020). Indeed, thanks to the substantially lower metabolic demand compared to traditional cycling (e.g. ∼15% lower rate of O_2_ consumption (V̇O_2_)) (Alessio et al., 2021; Berntsen et al., 2017; Bourne et al., 2020; Hoj et al., 2018; McVicar et al., 2022; Stenner et al., 2020) E-biking is accessible to a wider population and tolerable for longer commuting distances (Castro et al., 2019).

In this context, studies have reported that despite the lower metabolic demand, based on current physical activity guidelines, E-biking could still be considered a “moderate” activity that could aid individuals in meeting current recommendations (Berntsen et al., 2017; Bourne et al., 2018; Jenkins et al., 2023; McVicar et al., 2022). However, whether this presumption holds true remains uncertain because the validity of the guidelines-based methods used by these studies to characterize intensity of E-biking has been highly questioned (Iannetta et al., 2020, 2021; Jamnick et al., 2020). In this context, several recent studies have observed substantial discrepancies in terms of classification of exertional intensity between guidelines- [e.g., METs, percent of maximum heart rate (HR_max_), etc.] and metabolic threshold-based methods (Anselmi et al., 2021; Iannetta et al., 2020; Jamnick et al., 2020). Such discrepancy arises because guidelines-based methods do not consider the positions of the metabolic thresholds, which are highly individual in terms of both absolute and relative metabolic rates (Iannetta et al., 2020; Jamnick et al., 2020). Given that the magnitude of the metabolic stimulus in response to physical activity is function of the position of these thresholds and, therefore, of the metabolic *domains* that they define (Iannetta et al., 2020; Jamnick et al., 2020), using the current guidelines-based methods to quantify/classify intensity of E-biking may incur in erroneous interpretations of the associated metabolic demand. This is important to consider because small variations in work rate within and between intensity domains can significantly influence the metabolic stimulus of aerobic exercise (Brownstein et al., 2022; Iannetta et al., 2022) and, thus, its expected health benefits (Inglis et al., 2024; Ross et al., 2015). Thus, an evaluation of E-bike riding with respect to the threshold-based model is warranted to investigate whether such mode of transportation can be recommended to meet current physical activity recommendations.

Therefore, within a large and heterogeneous sample of individuals, this study aimed to quantify the reduction in metabolic demand associated with E-bike riding versus traditional cycling while considering the position of the gas exchange threshold (GET), which is the individual metabolic boundary portioning the moderate-from the heavy-intensity domain within the domain-based model (Iannetta et al., 2020; Jamnick et al., 2020). We hypothesized that the magnitude of the reduction in power output (PO) when riding an E-bike compared to traditional cycling is such that it will cause a shift in the domain of intensity according to the domain-based model of classification.

## Methods

For this study, 44 subjects (22 females) of different activity levels were recruited and completed the experimental procedures. The study was approved by the loal ethics committee (NP5282) and complied with the latest version of the Declaration of Helsinki. All participants signed an informed consent to participate in the study. None of participants had previous history of cardiovascular or metabolic disease nor were taking medications that could alter their cardiorespiratory responses to exercise.

### Exercise protocols

A schematic representation of the research design is depicted in **Figure 1**. Within two separate days, participants completed two experimental visits. The first one included performance of a ramp-incremental test in an environmentally-controlled laboratory to measure V̇O_2max_, GET, and the respiratory compensation point (RCP). The second visit was outdoor wherein participants completed twice a pre-determined course on the same bicycle equipped with an electrical motor, first without and then with electrical assistance. All sessions were completed at the same time of the day (±30 min).

**Figure 1.**
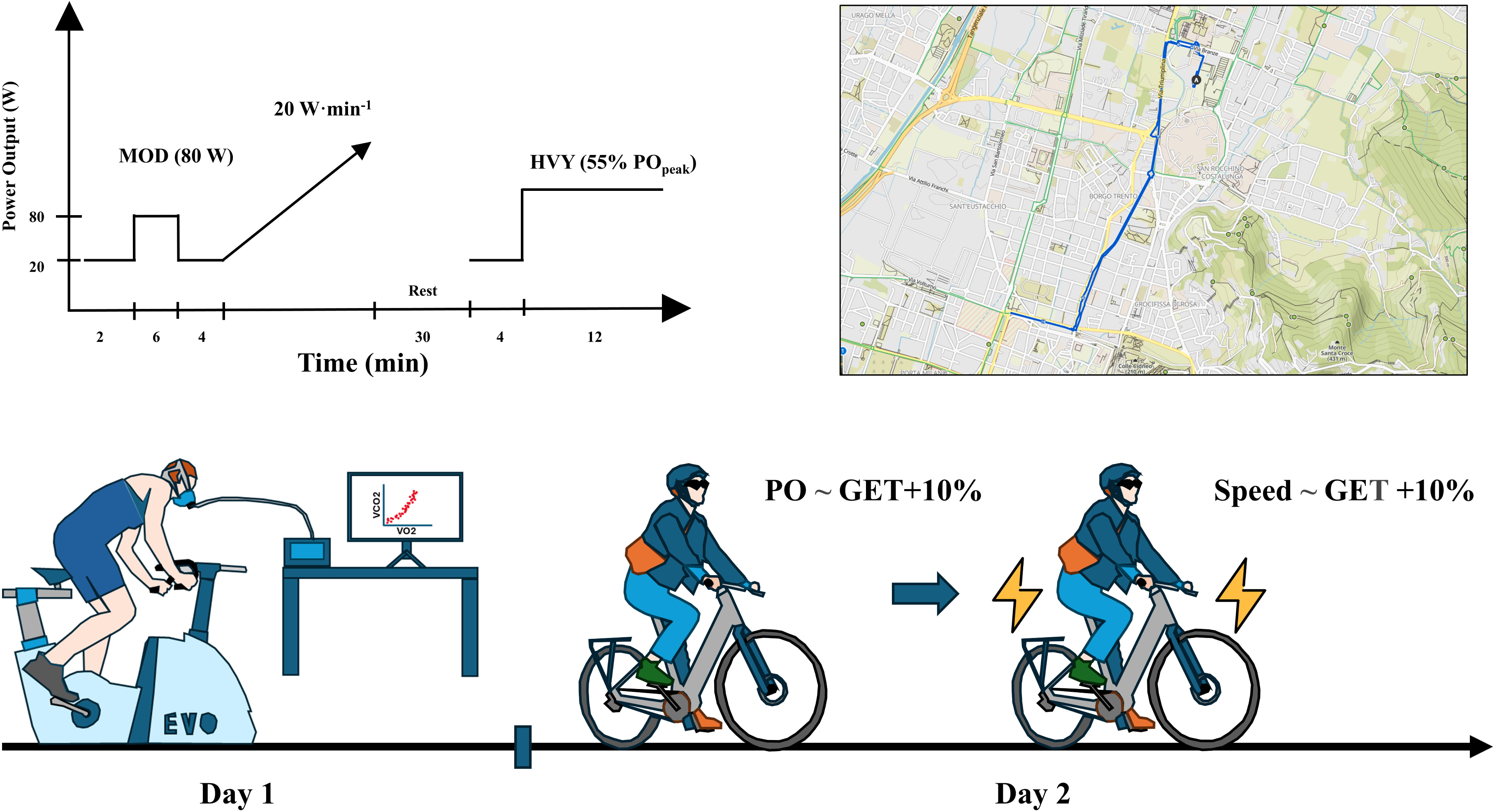
A schematic representation of the experimental procedures.

#### Ramp-incremental protocol

The ramp-incremental protocol consisted of one moderate-intensity step transition, which was followed by the ramped portion of the test, which in turn was followed by performance of a heavy-intensity step transition. The moderate- and the heavy-intensity step transitions served the purpose of retrieving the PO at GET and RCP, as previously described (Iannetta et al., 2023). The moderate-intensity step transition was performed at 80 W and lasted 6 min. Thereafter, following a baseline cycling of 20 W for 4 min, the ramp-incremental test began and was set at a rate of 20 W·min^-1^. Participants were instructed to exert maximally, and the test was stopped when they reached volitional exhaustion or when they could no longer maintain the targeted cadence for more than 10 s. Following ∼30 min of recovery, participants performed the heavy-intensity step transition, which was set at ∼55% of the peak PO (PO_PEAK_) obtained during the ramp-incremental test. During the entire protocol, participants were asked to maintain a cadence between 70 and 80 revolutions per minute. Gas exchange and ventilatory variables were monitored with a metabolic cart (K5, Cosmed, Rome, Italy), while heart rate (HR) was measured with a chest strap (HRM Pro Plus, Garmin, Schaffhausen, CH). The exercise protocol was completed on a computer-controlled bike ergometer (EVO, Monark, Sweden) which was also equipped with pedals capable of independently measuring PO (Rally Xc200, Garmin, Schaffhausen, CH). These same pedals were used to monitor PO during the outdoor session (see below).

#### E-bike outdoor rides

During the second experimental visit, participants rode an E-bike (Nuvola, Italwin, Bologna, Italy). The first ride was completed without electrical assistance, while the second ride was completed with electrical assistance turned on and set to the lowest level. The pre-determined course consisted of a 5.9-km long pathway, which took between 18 and 36 minutes to complete, depending on the fitness level of each participant. These durations are common for cycling commuting trips (Schantz, 2017; Schantz et al., 2020). The same power meter pedals used during the first laboratory session were also used to monitor and record PO during these outdoor rides.

During the first ride, participants were instructed to maintain an average PO ∼10% above GET identified during the first laboratory session. We selected this intensity for two reasons. Firstly, considering that GET occurs on average at ∼60% of V̇O_2max_ (Iannetta et al., 2021) and that the average V̇O_2_ during cycling commutes is reportedly ∼65% of V̇O_2max_ (Schantz et al., 2020), this choice ensured an individualized yet generalizable intensity of exertion. Secondly, as recently demonstrated (Inglis et al., 2024), an intensity just above GET is the minimum level of exertion recommended to elicit health benefits in individuals who have limited time to exercise.

After the first ride, a ∼20-min rest period was given to all participants to ensure that all physiological parameters would return to baseline. After this recovery phase, participants began the second ride, which was performed at the same average speed of the previous ride without electrical assistance. An investigator closely monitored each participant to ensure that they would follow the established speed. A smartwatch (Forerunner 945, Garmin, Schaffhausen, CH) was mounted on the bike’s handlebar to record PO, speed, and HR.

### Data Analyses

#### Identification of GET and RCP

To identify the GET, the relationship between V̇CO_2_ and V̇O_2_ (V-slope method) was plotted, and the point at which a change of slope in this relationship occurred was identified as the V̇O_2_ at GET (Beaver et al., 1986). This point was then corroborated by the identification of both the first change in slope of the relationship between V̇O_2_–minute ventilation (V̇_E_) and the concurrently beginning of the flattening (isocapnic buffering) of the relationship between the end-tidal CO_2_ pressure (P_ET_CO_2_) and V̇O_2_. RCP was identified as the point when P_ET_CO_2_ began a precipitous fall with respect to V̇O_2_ after the period of isocapnic buffering, which was corroborated by the coincidental second change in slope between V̇_E_-V̇O_2_ and V̇_E_/V̇CO_2_-V̇O_2_ relationships (Keir et al., 2022). Two expert investigators examined the plots and the average of the two estimates was used for subsequent analyses. If the difference in these estimates exceeded 100 mL·min^-1^, the two investigators would discuss the position of GET and RCP until a consensus was reached. To account for the V̇O_2_ kinetic dynamics at ramp-onset (normally referred as “mean response time”) which offsets the ramp-V̇O_2_ from its corresponding steady state-PO equivalent, the V̇O_2_ response was “left-shifted”, as per procedures outlined elsewhere (Iannetta et al., 2019). Briefly, the V̇O_2_ at 80 W of the moderate-intensity step transition was superimposed onto the ramp-derived V̇O_2_-PO relationship. The difference between the ramp-derived PO corresponding to this V̇O_2_ and the moderate-intensity PO (i.e., 80 W) was then calculated and used to “left-shift” the V̇O_2_ response to align it to its steady state equivalent. In a similar manner, to identify the PO corresponding to RCP, the relationship between the known V̇O_2_ and POs at GET and at the heavy-intensity step transition was extrapolated until the V̇O_2_ at RCP. The PO at the RCP was then retrieved by linear interpolation to the x-axis (Mackie et al., 2024).

#### Physiological responses during the outdoor rides

PO and HR collected during the outdoor rides were analysed using MATLAB (MATLAB, The MathWorks, Inc, Natick, MA, USA). The averages for PO and HR corresponded to the average values of the entire rides. To derive the metabolic equivalents (METs) associated with the rides performed without and with electrical assistance, for each individual the laboratory-derived moderate-and heavy-intensity V̇O_2_-PO relationships were used. For instance, if the average heavy-intensity PO of the ride without electrical assistance was 120 W, the corresponding V̇O_2_ was retrieved by linear interpolation of the slope of the relationship between the known V̇O_2_ and PO coordinates at GET and this PO (i.e., 120 W). The derived V̇O_2_ was then normalized by body mass and divided by 3.5, which is the nominal MET value (expressed in mL·kg^-1^·min^-1^) at rest (Jetté et al., 1990).

### Statistical analyses

Data are presented as mean ± standard deviation (SD). The Shapiro-Wilk test was used to assess the normality of the data collected during the lab test. An independent samples t-test was used to compare potential sex differences for the lab test variables where the data were normally distributed. If data were not normally distributed, the Mann-Whitney U test was employed. To investigate the effects of condition (without and with assistance) and sex, a repeated measures ANOVA was conducted, with assistance as a within-subjects factor and sex as a between-subjects factor. Mauchly’s test was used to assess the assumption of sphericity. In cases of non-sphericity and heterogeneity of variance, a repeated measures ANOVA with bootstrapping was performed (Johnston & Faulkner, 2021). Where significant main effects or interactions were found, Bonferroni post-hoc tests were conducted to examine pairwise comparisons between condition levels. All statistical analyses were performed using R (v. 4.3, R Core Team, 2023). Statistical significance was set at *P* ≤ 0.05.

## Results

### Participant characteristics and ramp-derived results

Participant characteristics and physiological responses obtained during the ramp-incremental test are presented in **Table 1**. On average, females had a lower PO_PEAK_ (*P*<0.001), V̇O_2max_ (*P*<0.001), GET (*P*<0.001) and RCP (*P<*0.001) expressed as both absolute and relative terms.

**Table 1.** Participants characteristics and data collected as part of the ramp-incremental test.

|  | Total = 44 | Males = 22 | Females = 22 | *P* value |
| --- | --- | --- | --- | --- |
| Age, years | 32±10 | 31±12 | 32±8 | 0.328 |
| Height, cm | 173±8 | 178±7 | 168±6 | <0.001 |
| Body mass, kg | 67±13 | 76±12 | 57±7 | <0.001 |
| PO<sub>PEAK</sub>, W | 242±68 | 291±56 | 193±37 | <0.001 |
| PO<sub>PEAK</sub>, W·kg<sup>-1</sup> | 3.6±0.7 | 3.8±0.8 | 3.3±0.5 | 0.015 |
| V̇O<sub>2max</sub>, L·min<sup>-1</sup> | 3.03±0.98 | 3.75±0.71 | 2.3±0.62 | <0.001 |
| V̇O<sub>2max</sub>, mL·kg<sup>-1</sup>·min<sup>-1</sup> | 44.6±9.9 | 49.7±9.4 | 39.5±7.7 | <0.001 |
| GET, W | 112±34 | 133±31 | 91±22 | <0.001 |
| GET, W·kg<sup>-1</sup> | 1.6±0.3 | 1.7±0.4 | 1.5±0.3 | 0.125 |
| GET, L·min<sup>-1</sup> | 1.72±0.54 | 2.05±0.51 | 1.40±0.32 | <0.001 |
| GET, mL·kg<sup>-1</sup>·min<sup>-1</sup> | 25.7±6.3 | 27.3±7.6 | 24.0±4.3 | 0.141 |
| RCP, W | 155±62 | 194±60 | 116±35 | <0.001 |
| RCP, W·kg<sup>-1</sup> | 2.2±0.7 | 2.6±0.9 | 1.9±0.4 | 0.008 |
| RCP, L·min<sup>-1</sup> | 2.54±0.84 | 3.12±0.67 | 1.9±0.51 | <0.001 |
| RCP, mL·kg<sup>-1</sup>·min<sup>-1</sup> | 37.5±8.9 | 41.5±9.4 | 33.5±6.4 | 0.002 |

Data are showed as mean ± SD. Sexes were compared using an independent t test.
PO<sub>PEAK</sub>: peak power output; V̇O<sub>2max</sub>: maximal rate of oxygen consumption; GET: gas exchange threshold; RCP: Respiratory compensation point.

### Performance and physiological responses during the outdoor rides

**Figure 2A** and **2B** show the PO and the HR during the rides without and with electrical assistance. The PO determined for the outdoor ride without assistance and corresponding to GET +10% corresponded to 124±37 W. There was no difference between this pre-determined PO and the actual PO recorded during the ride (127±35 W; *P*=0.163). The HR elicited by this PO was 127±19 bpm and corresponded to 69±8% of HR_max_. Overall, participants completed the course in 25±4 min at an average speed of 13.9±2.1 km·h^-1^. Time required to complete the ride (25±3 min; *P*=0.143) and speed (14.2±1.9 km·h^-1^; *P*=0.074) were nearly identical during the second outdoor ride completed with electrical assistance. However, expectedly, the average PO was lower (41±27 W, *P*<0.001). The difference in PO measured at the pedals between the rides without and with electrical assistance was 86±19 W, which equates to a delta percent of 69±12%. HR throughout the ride with electrical assistance (97±15 bpm, 53±7% HR_max_) was also lower (−16±5% HR_max_) compared to the ride without electrical assistance (127±19 bpm, 69±8% HR_max_; *P*<0.001). On average, we calculated that the METs associated with the ride without electrical assistance were 8.3±1.7 METs, while the ones associated with the ride with electrical assistance were 4.2±1.3 METs (*P*<0.001).

**Figure 2.**
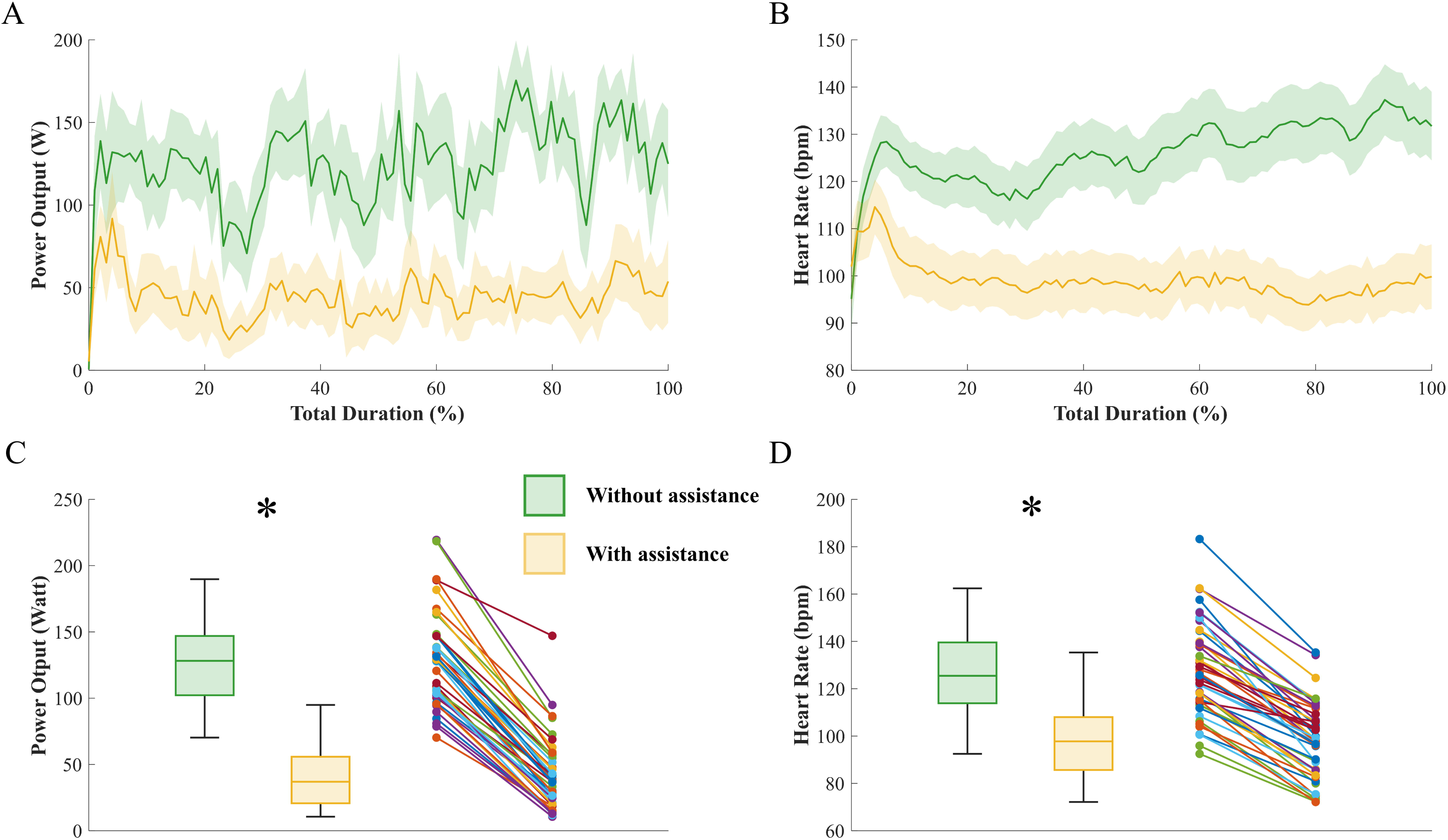
Mean ± 95% confidence intervals for power output (PO) (A) and heart rate (HR) (B) during rides with (orange) and without (green) electrical assistance. Boxplots and individual data (Panels C, D) display the distribution of participants for PO and HR.

### Sex-related differences

The average PO during the ride without electrical assistance was 150±33 W for males and 104±19 W for females (*P*<0.001). Despite this difference in PO between sexes, the speed (*P*= 0.161) and time (*P*= 0.126) to complete the ride without electrical assistance was similar (males 14.6±2.1 km·h^-1^; 24.5±4.0 min; females 13.3±1.8 km·h^-1^; 27.0±3.4 min). These were not different from the speed (*P*=0.279) and time (*P*=0.131) recorded during the ride with electrical assistance (males 14.8±1.8 km·h^-1^; 24.0±2.9 min; females 13.6±1.9 km·h^-1^; 26.5±3.6 min). When riding with electrical assistance, the PO was reduced to 54±30 W in males and to 28±15 W in females (*P*=0.001). This absolute reduction in PO was significant between sexes (males 95±21 W; females 76±10 W; *P*=0.001). Nonetheless, this reduced PO with electrical assistance corresponded in both sexes to the same percentage of the PO at GET (males 37±19%; females 31±11%; *P*=1.000).

During the ride without electrical assistance, HR was similar in males (122±20 bpm) compared to females (132±17 bpm; *P*=0.426). During the ride with assistance, HR was 92±15 bpm for males and 102±14 bpm for females (*P*=0.430). When normalized to HR_max_, during the ride without assistance, there was no difference between males (67±8% HR_max_) and females (73±7% HR_max_) (*P*=1.000). Similarly, during the ride with assistance, no difference between sexes was found (males 52±7% HR_max_, females 55±6 HR_max_; *P*=1.000). There was no difference between sexes in terms of METs during the ride without assistance (males: 8.8±2.0 METs; females: 7.8±2.3 METs; *P*=0.205) nor with assistance (males: 4.5±1.6 METs; females: 3.9±1.0 METs; *P*=1.000).

## Discussion

Considering the importance of an accurate characterization of exertional intensity (Iannetta et al., 2020, 2021) and given the shortcomings of the methods used by previous studies to classify intensity of E-biking (Alessio et al., 2021; Anderson et al., 2022; Bourne et al., 2018; Haufe et al., 2022; Hoj et al., 2018; Jenkins et al., 2023; McVicar et al., 2022), we investigated the metabolic responses of riding an E-bike compared to traditional cycling in relation to the exercise intensity domain schema. Findings were that riding a popular model of E-bike is associated with a shift of the domain of intensity (heavy→moderate), requiring a level of exertion that is substantially lower (∼36%) than the intensity associated with GET. Overall, these findings demonstrate that despite current guidelines would suggest E-bikes as suitable to meet current exercise intensity recommendations, when interpreted in the context of the exercise intensity domains and in relation to a specific intensity threshold (i.e., GET), the metabolic demand associated with routine bike commuting with electrical assistance is likely insufficient to evoke a metabolic stress that is high enough to promote substantial cardiorespiratory gains.

### Physiological responses

Compared to traditional cycling without electrical assistance, in this study riding an E-bike led to a reduction in PO of ∼86 W (or 69 %) and a reduction in HR of ∼30 bpm (or ∼23 %). Although these findings are in line with previous studies showing a reduction in metabolic demand when riding an E-bike, the extent of the reductions herein is somewhat larger than the one reported by previous studies. For example, Langford et al. (Langford et al., 2017) reported a 22 W decrease in PO and 5 bpm in HR between traditional cycling and E-biking. Similarly, Louis et al. (Louis et al., 2012) found a reduction of 13 W in PO and 3 bpm in HR when using a “moderate” assistance level at a speed of 16 km·h^-1^, while the reduction was ∼47 W and ∼13 bpm when using a “high” assistance level. Likewise, Simons et al. (Simons et al., 2009) observed reductions in the order of 17 W for PO and 7 bpm for HR as well as 24 W for PO and 11 bpm for HR when riding the E-bike with moderate and high levels of assistance, respectively. Finally, Sperlich et al. (Sperlich et al., 2012) found a reduction of 33 W and 28 bpm. Summarizing these and other findings through a meta-analytic approach, McVicar et al. (McVicar et al., 2022), calculated that the average reduction in PO and HR reported to date within the literature is in the order of 19 W for PO and 3.5 bpm for HR, respectively. As also mentioned herein, however, the variability amongst studies is quite large. Putative explanations for such variability may relate to the characteristics of the E-bike used in terms of the levels of assistance offered, the course selected (open roads vs cycling dedicated pathways), and the level of fitness (trained vs untrained). Furthermore, in some studies participants were allowed to choose their preferred speed during the ride with the E-bike which might have jeopardized the exact computation of the difference between riding with and without electrical assistance.

#### Sex Differences

Predictably, there were differences in PO between sexes for the rides without and with assistance. These differences can be attributed to the different levels of cardiorespiratory capacity between the two sexes (i.e., PO_PEAK_, males: 291±56 W *vs* females: 193±37 W; GET, males: 133±31 W *vs* females 91±22 W; *P*<0.001). While both sexes showed a decrease in PO and HR, males had a greater absolute value of PO in both conditions (*P*<0.001). However, when we adjusted PO for each individual’s GET, there was no difference between sexes. This means that relative to their individual capacity, males and females experienced a similar decrease in PO and comparable metabolic stress during the rides. This was further corroborated by the finding that neither the absolute nor relative reductions in HR differed between sex.

Despite males having higher PO than females, both sexes averaged similar speeds during the rides. This could be because males, who were taller and heavier (thus having a greater surface area), are likely less aerodynamically efficient. This lower aerodynamic efficiency would have required them to generate more power to maintain the same speed as females. This interpretation highlights the fact that sex is a variable that could modulate the effect of the electrical assistance on the metabolic demand of e-biking.

#### Implications of the findings

In recent years, the rapid spread of E-bikes has represented an opportunity for many to engage in more physically active behaviour and to contribute to the collective effort to reduce environmental pollutants (Johansson et al., 2017; Li et al., 2023; Sundfør et al., 2020). Whether, though, E-bikes can be used to induce substantial health gains has remained controversial. For example, although the majority of studies have shown some positive health benefits, such as in terms of glucose and blood pressure control (Alessio et al., 2024), when investigated under the lamp of cardio-respiratory fitness gains, the benefits of E-bike riding has remained discordant. For example, in two different studies, De Geus et al. (de Geus et al., 2013) and McVicar et al. (McVicar et al., 2023) found no changes in V̇O_2max_ after six weeks of E-bike riding, while Peterman et al. (Peterman et al., 2016) showed a significant increase after 4 weeks. The reason for such discrepancies may lie on the fact that, for many, the use of E-bikes is associated with a metabolic demand that is too low to stimulate changes in cardiorespiratory fitness. In this context, a recent study has demonstrated that in order to maximize the chances of positive cardiorespiratory benefits, previously inactive individuals are required to exceed intensity associated with the first metabolic threshold (GET) (Inglis et al., 2024; Ross et al., 2015). From Figure 3, it is possible to appreciate how, regardless of the fitness level of the individuals, routine bike commuting with electrical assistance unlikely allows individuals to surpass this threshold. Of course, it cannot be excluded that with larger volumes of E-bike riding exceeding current physical activity recommendations (>150 min/week), this reduced intensity may still suffice to induce changes in cardiorespiratory fitness (Peterman et al., 2016). However, this is unlikely to occur in the context of regular daily bike commuting.

**Figure 3.**
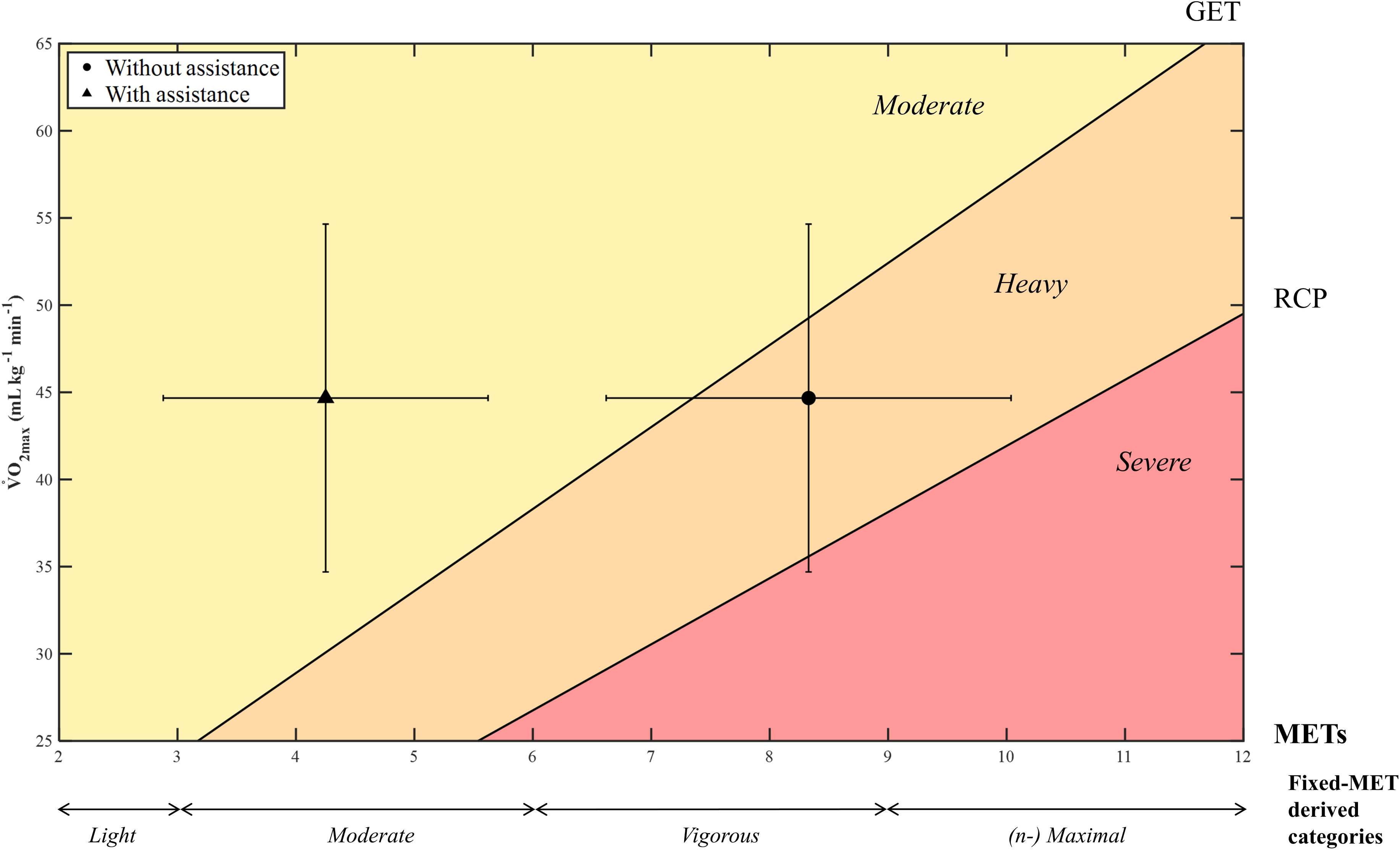
A schematic representation of the progression of the exercise intensity domains as function of V̇O_2max_ as measured in the present dataset. The different colored areas depict the three intensity domains: moderate (yellow), heavy (orange), and severe (red) in relation to fitness level (i.e., V̇O_2max_) (y-axis). The bottom portion shows the four conventional MET-derived categories of intensity. The circle within the graph represents the average (±SD) METs computed without electrical assistance, while the triangle represents the average METs (±SD) observed during the ride with electrical assistance. This schematic underscores that, although the METs of the ride with electrical assistance fall within the moderate-intensity range according to the traditional METs-based classification, if interpreted according to the intensity domain schema, these METs values are far below the METs associated with the GET and the minimum exertional intensity associated with cardiorespiratory benefits (Inglis et al., 2024).

Considering all this, although the metabolic demand of E-bike falls inexorably within the fixed-MET “moderate” intensity category (Figure 3), when contextualized within the concept of intensity domains, such finding questions the ability of current guidelines-based methods to capture with accuracy the demand of E-bike riding. Indeed the fact that on average riding with electrical assistance is associated with an intensity of ∼4.5 METs (Figure 3; see also (Sperlich et al., 2012))may give a misleading impression that it is suitable to satisfy the minimum intensity required to improve cardio-respiratory fitness. One needs to consider, however, that exercise intensity remains a relative concept, and the variable distance between the intensity associated with E-Biking and the intensity associated with the first metabolic threshold (i.e., GET) questions not only the use of the guidelines-based methods of classification but also the consequential proposition of that E-Biking can be used to improve cardio-respiratory fitness. Importantly, we do not advocate against the use of E-bikes. As mentioned before, there are many health- and environmental-related circumstances in which E-bikes can be extremely useful and should be promoted (Johansson et al., 2017; Li et al., 2023; Sundfør et al., 2020). Our data, though, urge a reconsideration of how intensity of E-bike is quantified by and interpreted within current physical activity guidelines. Considering our results, in the future, E-bike motor developers should aim to engineer systems capable of providing levels of assistance that are individualized to the users’s level of fitness. In this manner, this would ensure that intensity of the physical activity remains within a range of intensity that promotes health benefits.

#### Methodological Considerations

This study is the first to evaluate the metabolic demand of e-biking relative to a specific individual metabolic threshold (i.e., GET). However, some methodological considerations need to be addressed. First, the E-bike used in the present study is a popular model that is widely available in many municipalities and activated when the user begins to pedal. This specific model provides a fixed amount of assistance which we set at the minimum level. It is likely that the magnitude of the reduction in metabolic demand would have differed had we used a different E-bike model. Our choice to select a publicly available E-bike model is justified by the need to ensure generalizable conditions. Secondly, the speed achieved during the ride without assistance was below the average speed of normal commutes (Schantz, 2017; Schantz et al., 2020). This was because the E-bike model used in our study is heavier (∼25kg) than traditional bikes. We reasoned, however, that it was important to ensure similar conditions between the assisted and un-assisted rides. Unlike previous studies that have somewhat jeopardized their findings by employing different bike models between the rides with and without assistance. In this context, measuring the PO exerted directly at the pedals gives us confidence regarding the accuracy of our results.

## Conclusion

There are a number of health- and environmental-related advantages of using E-bikes for commuting (Bourne et al., 2018; Johansson et al., 2017; Li et al., 2023; Sundfør et al., 2020). Despite the fact that current intensity guidelines would predict that exertional intensity associated with E-bike riding is sufficient to elicit cardiorespiratory benefits in a group population of recreationally active individuals, when compared to the intensity domain schema, it is unlikely that this mode of transportation could achieve this goal. Therefore, the present study, while not intending to discourage the use of E-bikes, does suggest caution when interpreting exertional intensity associated with this mode of transportation.

## Acknowledgements

The results of the study are presented clearly and honestly without fabrication, falsification, or inappropriate data manipulation, and the statement that the results of the present study do not constitute endorsement by ACSM.

